# Structural and Mechanistic Usage and Preference of Human, Dog, and Bat Receptors by Rabies Virus

**DOI:** 10.1101/2023.10.28.564510

**Authors:** Manar E. Khalifa, Mustafa Atasoy, Mohammed Rohaim, Leonie Unterholzner, Muhammad Munir

## Abstract

Rabies is a lethal zoonotic viral disease causing approximately 59,000 human deaths annually. Recently, several cellular receptors for rabies virus (RABV) entry and internalization have been identified. However, none of these receptors have been demonstrated to be indispensable for RABV entry. Here we describe the RABV receptor preference *in vivo*, utilizing a replication-competent vesicular stomatitis virus (VSV), in which the VSV surface glycoprotein was replaced with rabies virus glycoprotein. To investigate the specific role of RABV receptors in promoting RABV entry in non-permissive cell line, HaCaT cells were used as a cellular model refractory for RABV infection. Employing virus binding and quantification studies, we demonstrated that ITGB1 and mGluR2 are potential receptors for RABV entry and replication. Consequently, knockout (KO) cell lines corresponding to each of the ITGB1 and mGluR2 receptors were generated using CRISPR/Cas9 mediated knockout. Surprisingly, RABV was still able to enter and replicate in the generated KO cell lines, yet the replication and entry of RABV in KO cells lacking mGluR2 and ITGB1 were significantly reduced; respectively. These findings suggest that RABV utilize these receptors in series rather than sequentially. To test whether RABV utilizes similar receptor preference among human, dog, and bats, the A549, Pa-Br and MDCK cell lines that overexpress receptor orthologs from their respective species were infected with rVSV-dG-RABV-G-GFP and quantified for virus binding and released virus progeny. Our findings revealed that in human cells, ITGB1 increased virus entry, while nAChR enhanced virus replication. In bat cells, ectopic expression of nAChR allowed enhanced virus entry and internalization. While MDCK cells overexpressing ITGB1 enhanced the levels of virus entry and replication. Conclusively, our study, reveals the RABV distinct receptor preference, influenced by the underlying pathways that occur during the interaction between the virus and receptor in different cell lines. Additionally, it emphasizes the significance of host-specific factors in virus entry and replication.

**Graphical Abstract:** 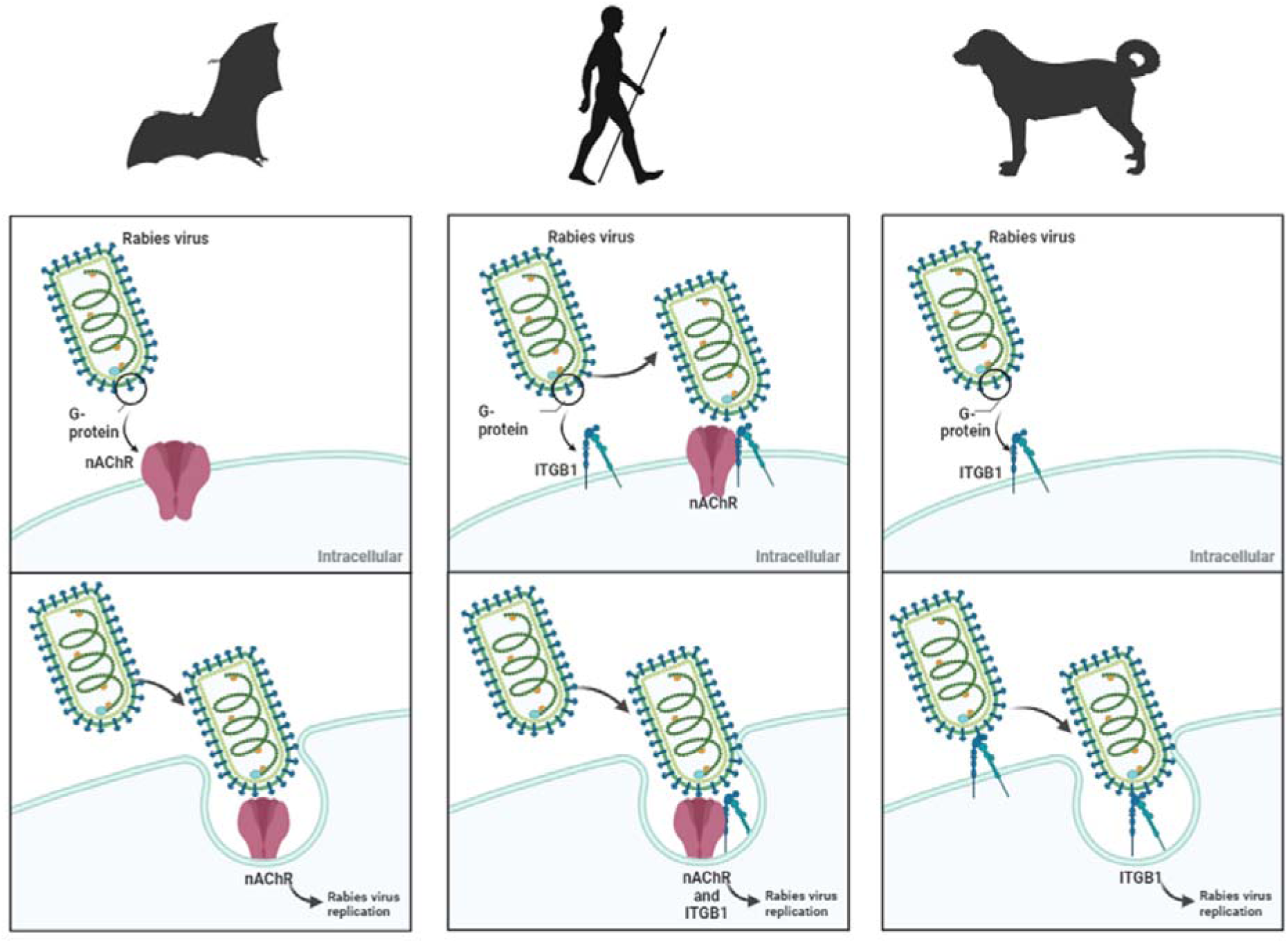

**Author Summary:** Rabies is a fatal neurological disease, characterized by broad host range and tissue tropism. In accordance with the global goal of eliminating dog-mediated human rabies by 2030, studying the underlying mechanism of RABV entry across distinct species would enable adjustment of RABV control strategies. Owing to RABV wide tropism, multiple cellular receptors have been identified for RABV entry into host cells. Previous studies have proposed that some of RABV receptors could serve as promising candidates for development of antiviral drugs (1). From this perspective, we focused on elucidating RABV receptor preference for viral entry in human, dog, and bat cells. In addition to determining whether RABV utilizes these receptors in parallel or in series which would indicate the potential of the identified RABV cellular receptors as targets for antiviral drugs against rabies. Our results demonstrated varying receptor preference of RABV across species. In addition to revealing that none of RABV receptors solely, govern the broad host range of rabies, suggesting that RABV antiviral drugs targeting host cellular factors may not effectively inhibit RABV entry into cells, while antiviral drugs targeting virus glycoprotein may exhibit greater efficacy. Collectively, our study, contribute to providing mechanistic model for RABV entry in different species.

## 1. Background

Rabies remains a fatal neglected tropical disease, despite imposing a public health significance by causing up to 60,000 human deaths annually with 30%-50% of these cases occurring in children according to the World Health Organization (WHO) (2). The causative pathogen, rabies virus (RABV), is a negative single stranded, enveloped RNA virus, which belongs to the family *Rhabdoviridae*, its genome is approximately 12 kb, encoding five structural proteins: nucleoprotein (N), phosphoprotein (P), matrix protein (M), glycoprotein (G), and an RNA-directed RNA polymerase (L) (3). Rabies G protein is exposed on the viral envelope, and it is the sole protein responsible for attachment with the cellular receptors (4). Upon attachment of G protein with cellular receptors, it triggers virus internalization to the cellular compartments *via* receptor mediated endocytosis (5). Despite the advancement in rabies research which identified multiple cellular receptors for RABV (1,6–10), none of the previous studies have identified any of these receptors as indispensable for virus entry. To date, six cellular receptors have been identified to facilitate RABV entry such as nicotinic acetylcholine receptor α1 (nAChRα1) (11), neural cell adhesion molecule, (NCAM) (9), metabotropic glutamate receptor 2 (mGluR2) (7), integrin beta 1 (ITGB1) (1) and transferrin receptor protein 1 (10).

Rabies virus represents a historically broad host and cellular range with the ability to infect diverse cell types among multiple species (12). Globally, dogs are associated with the human rabies cases in Africa and Asia, which has consequently linked dogs to RABV spillover events (13). Despite the circulation of RABV for centuries in domestics dogs, bats have been recently identified to be RABV ancestral reservoirs (14). Given that canine rabies is mostly eliminated in the Americas, bat born rabies is extensively emerging especially through the common vampire bats. The most common species contributing to bat rabies is *Desmodus rodontus* which are prevalent in South America to North Mexico (15). Multiple factors contribute to the substantial risk imposed by the bats acting as RABV reservoirs. One of the most crucial factors is their complete dependence on blood to survive which allows virus transmission to their prey through feeding (16). This results in primary and secondary spillover events, for example, RABV could be transmitted from the *D. rotundus* to cats (primary spillover) and from cats to humans (secondary spillover) (17).

Sine only microbats have been reported to act as RABV reservoirs and none of the previous surveillance data have shown whether any of the bat species belonging to the megabats could serve as intermediate host for RABV. The *P.alecto* is a bat genus belongs to megabats and has been previously reported to be associated with Australian bat Lyssavirus infection (18). Thus, in this study, we investigated whether *P.alecto,* representing one of the mega bats could be susceptible to RABV infection or not.

In the current study to understand the RABV receptor preference and the underlying mechanism of RABV infection in human, dog, and bats, we used a replication competent virus in which we introduced RABV G open reading frame (ORF) in VSV genome with reverse genetics with a green fluorescent protein (GFP) as a marker of infection. Along with utilizing HaCaT cell line as refractory for RABV infection which lacks host factors necessary for RABV replication allowing the study of RABV receptor preference. Our receptor preference data showed that none of the previously identified known RABV receptors were capable of rendering HaCaT cells susceptible for RABV infection, yet ITGB1 and mGluR2 yielded the highest levels of virus binding and replication; respectively.

Consequently, we explored the entry and replication of rVSV-dG-RABV-G-GFP into ITGB1 and mGluR2 A549 KO cell lines. Strikingly, our results showed the ability of rVSV-dG-RABV-G-GFP virus to enter and replicate in KO cells. However, entry and replication of rVSV-dG-RABV-G-GFP were significantly reduced in ITGB1 and mGluR2 KO cell lines; respectively compared to wild-type A549 cells. To further demonstrate whether the G protein of dog related RABV would have different permissiveness spectrum among cell lines of human bats and dogs. We comparatively assessed the rVSV-dG-RABV-G-GFP replication pattern on Pa-Br, A549 and MDCK cells overexpressing RABV orthologs receptors. From these experiments we obtained two conclusions; our first conclusion is that to our knowledge we are the first to report that the Pa-BR (Pteropus alecto brain) cell line which is derived from the *P.alecto* brain could be infected with the RABV. Secondly. is that rVSV-dG-RABV-G-GFP displayed differential receptor preference among the tested cell lines representing each of human dog and bat. The obtained findings suggest that none of the known RABV receptors defines its host range and thereby the design of antivirals targeting RABV receptors would not be effective in disease control.

## 2. Methods

### 2.1 Cells and viruses

Human keratinocytes (HaCaT), Human embryonic kidney cells (HEK293), Human lung carcinoma cells (A549), Chicken embryonic fibroblast (CEF), chicken origin DF-1 cells (DF-1), and Canine Madin-Darby Canine kidney cell (MDCK) were obtained from ATCC and maintained in Dulbecco’s modified Eagle’s medium (DMEM) (high glucose, GlutaMAX Supplement, pyruvate) (Life Technologies), supplemented with 5% foetal bovine serum (FBS) and 1 X antibiotic antimycotic solution. The *P.alecto* brain cell line (Pa-Br) was maintained in DMEM (F-12/HAM) supplemented with 10 % FBS and 1 X antibiotic antimycotic solution.

All cell lines were incubated at 37 °C and 5% CO2. Upon reaching 80% confluency, culture medium was aspirated and washed once with PBS for infection. Infection studies were performed with rVSV-dG-RABV-G-GFP. For the preparation of rVSV-dG-RABV-G-GFP, a BlueScript backbone plasmid under T7 polymerase promoter control encoding VSV Indiana backbone in which the VSV glycoprotein (G) gene has been replaced with GFP (pVSV-ΔG-GFP) was purchased from Kerafast. The pVSV-ΔG-GFP was modified by cloning the codon optimized ORF of RABV G protein of Egyptian strain (GenBank accession number MK760770.1) via MluI and NheI restriction sites, yielding a pVSV-dG-RABV-G-GFP. For recovery of the infectious virus from pVSV-dG-RABV-G-GFP, simultaneous transfection of the modified plasmid pVSV-dG-RABV-G-GFP along with VSV helper plasmids was carried out as previously described (19). Briefly, BHK-21 cells were seeded in a 6 well plate for 70-90% confluency, in the next day, the growth medium was aspirated, cells were washed with PBS. Followed by infecting the cells with recombinant fowl pox virus (rFPV), as a source of T7 promoter for 2 hrs, then the inoculum was removed, and infected cells were washed 3 X with PBS. Subsequently, BHK-21 cells were co-transfected with pVSV-dG-RABV-G-GFP, pBS-N-ФT, pBS-P-ФT, pBS-L-ФT, and pBS-G-ФT Plasmids. The transfection mixture was prepared by mixing the plasmids with turbofect transfection reagent, diluted in 500 μL Opti-MEM and incubated at RT for 25 min, followed by dropwise addition of transfection mixture to the cells and incubation at 37 °C and 5% CO2 overnight. After 72 hrs, the plate was frozen and thawed 3 times for harvesting. The recovered virus was centrifuged at 300 x g, aliquoted and stored at -80 °C for future use.

Plasmids: Plasmids encoding the cDNAs encoding ITGB1, mGluR2, nAChR orthologs in addition to ORF of *P.alecto* NCAM and p75 were codon-optimized and synthesized by (Bio-Basic Inc.) and cloned into pCAGG vector with a C-terminal FLAG tag.

The pVSV-dG -GFP and VSV helper plasmids were purchased from Kerafast company. All plasmid maps and cloning primer sequences are available upon request.

### 2.2 Plaque assay

HaCaT, Pa-Br, A549 wild-type (WT), A549 KO and MDCK cells in a 24-well plate were transfected with the corresponding plasmid expressing receptor orthologs or with empty vector for 48 hrs. After 48 hrs, cells were infected with rVSV-dG-RABV-G-GFP at MOI = 5 at 4 °C for 1 h. The unbound virus was removed through washing 3 times with ice-cold PBS. Plates were then incubated at 37 °C for 30 hrs. After 30 hrs post infection, virus supernatant was collected, and 10-fold serial dilutions were prepared in serum free medium (DMEM). Virus dilutions were used to infect BHK-21 cells in 6-well plate. The Plates were incubated for 2 hrs at 37 °C with shaking every 20 minutes to ensure uniform distribution of the virus dilutions. Cells were washed twice with PBS following the removal of the virus inoculum and the cells were overlaid with 4 mL of the plaque overlay media as previously described (20). The Plates were kept at 37 °C for 72 hrs, then cells were fixed with 4% paraformaldehyde (in 1 X PBS) for 1 hr on a shaker at room temperature. The overlay medium was aspirated after fixing the plates and stained with 0.5% crystal violet for 1 hr. The plates were then washed with water and the number of formed plaques were counted and determined in plaque forming units (PFU).

### 2.3 Genetic Knockout using CRISPR/Cas9

The design of single guide RNA (sgRNA) targeting the fourth and second exons of the human ITGB1, and mGluR2; respectively, was carried out using the online Benchling web tool (https://benchling.com) to minimize the off-target cleavage. For the forward (FWD) gRNA oligo sequence, an overhang of the (CACCG) was added at the 5’ end. The extra G added after the restriction site to ensure efficient transcription initiation from the U6 promoter. While an overhang of (AAAC) was added at the 3’ end of the reverse complement of the gRNA oligo sequence. sgRNA sequences for ITGB1 disruption were as follows.

5’-CACCGAATCGCAAAACCAACTGCTG-3’,

5’-AAACCAGCAGTTGGTTTTGCGATT-3’

sgRNA sequences for mGluR2 disruption were as follows.

5’-CACCGGGTCGCATAAGAGCCGTCG-3’,

5’-AAACCGACGGCTCTTATGCGACCC-3’

The sgRNA for each of ITGB1 and mGluR2 were synthesized and ligated in pSpCas9(BB)-2A-Puro (PX459) V 2.0 vector (ADDGENE, UK) using BbsI restriction site. For the generation of stable A549 knockout cell line, cells were seeded in 6 well plate and the next day cells were transfected with the recombinant Cas9 plasmids with either ITGB1, or mGluR2 gene-specific sgRNA using lipofectamine. The cells were incubated for 24 h prior to addition of the selective antibiotic medium containing the puromycin (Gibco, China) at a concentration of 2 µg/mL, after addition of the antibiotic selection medium, the growth media was replaced every 48 hrs for removal of non-transfected cells. Followed by isolation of cell clones w in 96 well plates for obtaining single cell clones using limiting dilution technique. The single-cell clones were visible by microscopy within 10-14 days, which were expanded and transferred into a larger plate until enough cells were obtained for harvesting. Single cells were further screened using PCR and sequence analysis to detect and characterize the indels in the gene of interest (21). The sanger sequences were analysed using inference of CRISPR Edits (ICE) (22). Positive cell clones were screened by genomic PCR using primers flanking the target exons. Genotyping primer sequences for ITGB1 KO were as follows:

FWD: 5’-CAATTTTCATTTATACCTATATTTTATATGTCA-3’,

REV: 5’-AATTAATACTTTCTGAATCTTTAACAAAATTTACTTTGAA-3’

Genotyping primer sequences for mGluR2 KO were as follows:

FWD: 5’-CAGAAGGGCGGCCCAGCAGAGGACTGTGGTCCTGTCAATGAG-3’

REV: 5’-AAATCGACCACCATTATGTGACCAGGGCACTTTCTTAGCTTC-3’.

### 2.4 Western blotting

Expression of cellular receptors in pCAGG-FLAG plasmids was confirmed through western blot analysis using rabbit anti-FLAG as primary antibody. The HEK293, or A549 or MDCK cells were transfected with plasmids encoding the corresponding receptor orthologue. After 36 hrs post transfection, cells were washed with ice cold PBS and trypsinized. Followed by cell pelleting, and resuspension of cell pellet with 100 μL ice cold lysis buffer composed of NP-40 lysis buffer with protease inhibitors (Thermo Scientific, USA) and kept on ice rocker for 30 minutes for whole cell lysate harvest. For removal of cell pellet, the mixture was centrifuged at 14,000 x g for 5 minutes. Cell lysates were then mixed with 2 x of the NuPAGE™ LDS Sample Buffer 4 x (Novex, Life Technologies, USA) with addition of 10% β-mercapto-ethanol and incubated for 5 min at 98 °C. Samples were then electrophoresed in polyacrylamide gels and transferred into PVDF membrane (Thermo Scientific™), for blotting. Upon membrane blotting, the membrane was blocked in 5% non-fat dry milk in PBST (0.5% tween-20 in PBS) for 1 hour. Afterwards, the membrane was washed once with (0.5% tween 20 in PBS). Consequently, the membrane was incubated with anti-FLAG polyclonal primary antibody produced in rabbit (Sigma-Aldrich Cat# F7425, RRID: AB_439687) at 4° C overnight Then the membrane was washed 3 times in PBS-T (0.5% tween 20 in 1 x PBS), 5 min/wash. Followed by incubation with Goat Anti-Rabbit IgG - H&L Polyclonal antibody, HRP-Conjugated (Abcam Cat# ab6721, RRID: AB_955447) for 2 hours at room temperature. Thereafter, the membrane was washed 3 times with PBST (0.5% tween 20 in PBS). Eventually, the membrane was incubated for 1 minute with Pierce ECL Western Blotting Substrate (Thermo Fisher) added 1:1 of detection reagents 1 and 2. Membranes were visualized using ChemiDoc™ MP imaging System (Bio-Rad Chemidoc, Hercules, CA, USA). For loading controls, identical protein lysate aliquots were incubated with mouse monoclonal anti-alpha tubulin antibody ((Abcam Cat# ab15246, RRID: AB_301787) with secondary antibody rabbit anti-Mouse IgG H&L (Abcam Cat# ab6721, RRID: AB_955447) Abcam) (23).

### 2.5 Immunofluorescence assay

HEK293, A549 and MDCK Cells were cultured on cover slips, upon reaching 70% confluency, cells were transfected with plasmids expressing receptor orthologs. Twenty-four hrs post transfection, cells were washed once with 200 μL PBS, then fixed for 1 hour with 200 μL 4% PFA at room temperature, then washed with PBS. Afterwards, cells were treated with 0.1% Triton X-100 (Sigma-Aldrich, T8787) in PBS for 10 minutes, then cells were washed once with PBS and blocked with 0.5% bovine serum albumin (BSA) (Sigma-Aldrich, 05482) in PBS for 1 hour. After blocking, BSA was aspirated, and cells were probed with anti-FLAG polyclonal primary antibody produced in rabbit (Sigma-Aldrich Cat# F7425, RRID: AB_439687) overnight at 4° C. Next day, cells were washed 3 x with PBS for 5 min. each, followed by incubating the cells with Goat Anti-Rabbit IgG (H+L), Alexa Fluor 488 Conjugated antibody (Molecular Probes Cat# A-11008, RRID:AB_143165) for 1 hour at room temperature, followed by washing cells 3 x with PBS, 5 min/wash, and a final wash with distilled water was carried out. Thereafter, the nuclei were stained with 4’,6-diamidino-2-phenylindole (DAPI) (Thermo Fischer, 62247) for 30 minutes (1:10,000). After nuclei staining, coverslips were prepared for mounting over microscopic slides using VECTASHIELD anti fade medium (Vector Laboratories, ZH1108) and fixed with clear nail varnish. Images were acquired with laser confocal microscope, (LSM880). Analysis and data processing were executed using Zeiss software (24).

### 2.6 Virus binding assays flow cytometric analysis (FC)

HaCaT, Pa-Br, A549 WT, A549 KO and MDCK cells in a 24-well plate were transfected with the corresponding plasmid expressing receptor orthologs or with empty vector for 48 hrs. After 48 hrs, cells were infected with rVSV-dG-RABV-G-GFP at MOI = 5 at 4 °C for 1 h. the unbound virus was removed through washing 3 times with ice-cold PBS. Subsequently, cells were collected and sustained with live-dead marker (Thermo Fisher, USA), followed by washing and permeabilization with 1x permeabilization buffer, then washed and resuspended in PBS with 2% FBS for 30 min for blocking. Upon blocking, cells were surface-stained with a rabbit polyclonal anti-FLAG antibody (Sigma-Aldrich Cat# F7425, RRID: AB_439687) and mouse RABV-G antibody ((Bio-Rad Cat# MCA2828, RRID: AB_1125351) kept for 1 hr in cold room. Then cells were washed 3 times with FACS buffer (1X PBS supplemented with 2% FBS) followed by staining with corresponding secondary antibodies goat α-rabbit IgG Alexa568 (Molecular Probes Cat# A-11011, RRID:AB_143157) and goat anti-mouse IgG Alexa-Fluor 488 ((Thermo Fisher Scientific Cat# A-11001, RRID:AB_2534069) for 45 min in FC buffer, on ice to analyse expression levels and RABV-G receptor engagement, respectively. The concentration of the antibodies was used at 1.5 µg/mL after secondary antibody staining, cells were washed three times, followed by resuspension in FC buffer to be analysed using CytoFLEX Flow Cytometer (Beckman Coulter, USA).

### 2.7 Virus infection assays flow cytometric analysis

Pa-Br, A549 WT, A549 KO and MDCK cells in a 24-well plate were transfected with the corresponding plasmid expressing receptor orthologs or with empty vector for 48 hrs. After 48 hrs, cells were infected with rVSV-dG-RABV-G-GFP at MOI = 5 at 4 °C for 1 h. the unbound virus was removed through washing 3 times with ice-cold PBS. Plates were then incubated at 37 °C for 30 hrs. Subsequently, cells were collected and stained with live-dead marker (Thermo Fisher), followed by cell pelting, washing and permeabilization with 1 x permeabilization buffer. After 15 min, cells were washed and resuspended in FC buffer for 30 min for blocking. Upon blocking, cells were surface stained with a rabbit anti-FLAG IgG antibody kept for 1 hr in cold room. Then cells were washed 3 times with FC buffer followed by staining with goat α-rabbit IgG Alexa-568 (Molecular Probes Cat# A-11011, RRID: AB_143157); for 45 min in FC buffer to analyse the GFP percentage corresponding to virus infection on transfected cells. The concentration of the antibodies was used at 1.5 µg/mL after secondary antibody staining, cells were washed three times, followed by resuspension in FACS buffer to be analysed using CytoFLEX Flow Cytometer (Beckman Coulter, USA).

### 2.8 Gating strategy for the transfected/infected cells and entry experiments

The analysis of flow cytometry data was conducted using FCS Express 7 (DeNovo Software, Los Angeles California). Cells were gated, against the PB450-A channel, forward and side scatters to allow the selection of the live, singlet cell populations. The percentage of RFP positive cells populations corresponding to cells expressing the receptors was calculated. From the RFP positive cell populations, the GFP positive cells were gated. For the entry experiments, the GFP positive was referred to as entry GFP % of transfected cells for the infection experiments, the GFP positive cell populations were referred to as GFP% of transfected cells. The compensations of the samples were carried based on using the following controls: negative cell control (uninfected, non-transfected), FITC positive sample only (infected only; GFP positive only) and RFP positive sample only (transfected only; RFP positive only). Upon compensation, data were plotted in representative histograms, along with empty vector (EV) control which served as infected, transfected empty vector control as well as the cell control (uninfected, non-transfected cells) which served as negative control to regulate the gating of the GFP percentage.

### 2.9 Statistical analysis

The statistical analysis was conducted using GraphPad Prism 9. The significance testing was carried out using a student t-test, in case only two groups were compared. Experimental means were compared using one-way ANOVA (analysis of variance) when multiple comparisons were required for a single factor. The presented data usually represents three biological replicates with standards error of means (SEM). *P* values less than 0.05 were regarded as statistically significant as indicated: ns: non-significant; p>0.05, *p < 0.05, **p < 0.01, ***p < 0.001, ****p < 0.0001.

## 3. Results

### 3.1 HaCaT cell line has been identified as refractory for rVSV-dG-RABV-G-GFP infection

Using reverse genetics system, we developed a replication competent vesicular stomatitis virus (rcVSV) in which the VSV G was swapped with a reporter GFP gene along with the insertion of the rabies surface glycoprotein to allow studying the RABV tropism. The infection with the generated rVSV-dG-RABV-G-GFP resulted in GFP expression on infected cells. A wide range of cell lines were tested for their susceptibility to rVSV-dG-RABV-G-GFP, all the tested cell lines showed susceptibility to infection **(Fig S1).** Only the immortalized human keratinocyte cells (HaCaT cells) showed no GFP upon infection with the rVSV-dG-RABV-G-GFP **(Fig 1 A)**.

**Figure 1.**
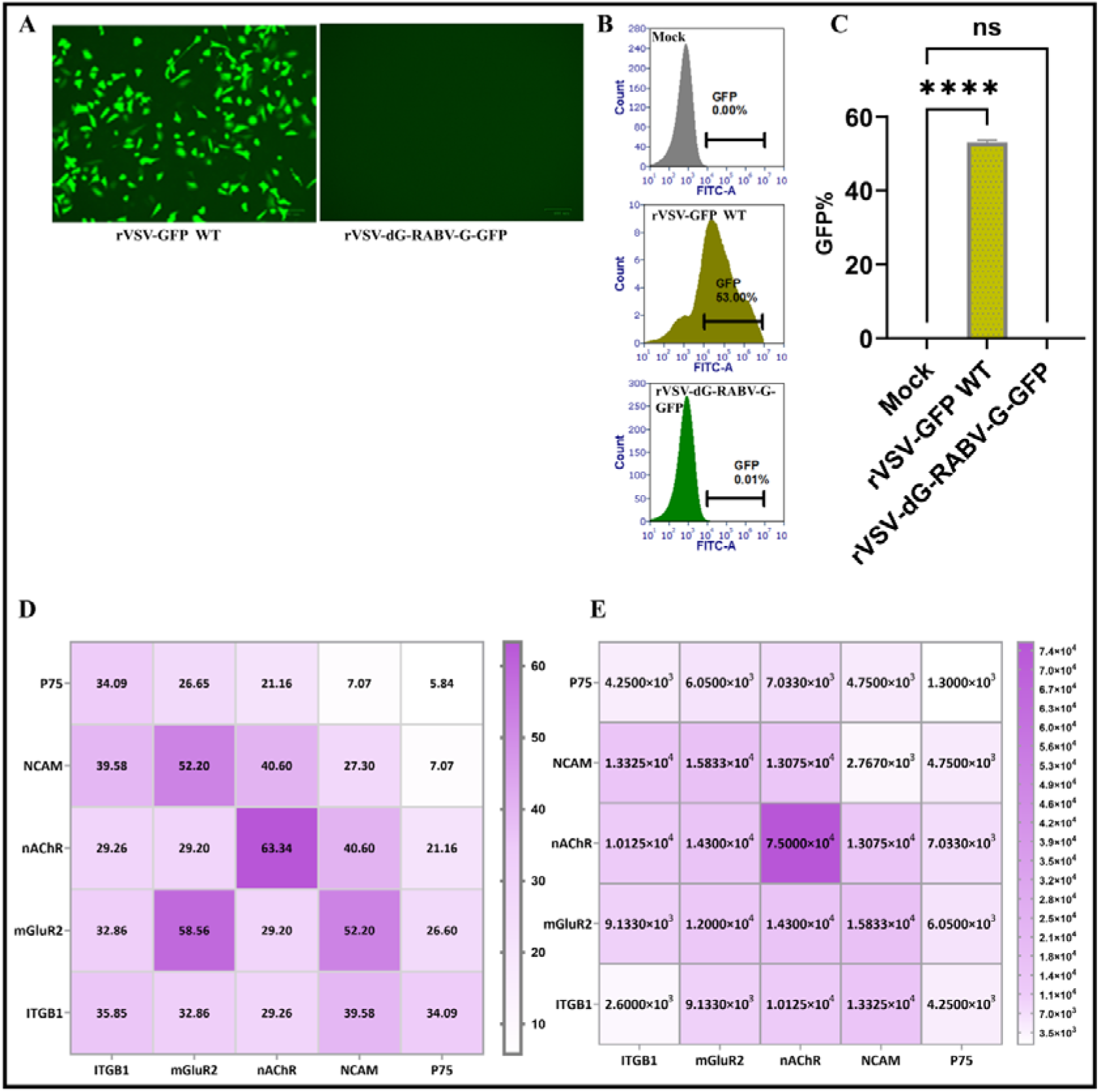
HaCaT cells are permissive to rVSV-GFP-WT virus infection. but not VSV-dG-RABV-G-GFP. ***(A).*** *Representative microscopic green fluorescence images of HaCaT cells infected with VSV-WT (left) and rVSV-dG-RABV-G-GFP (right), scale bars size, 100 µm **(B).** Representative histograms showing the percentage of infected cells as quantified by flow cytometry at 30 hpi with HaCaT cell control, HaCat cells infected with VSV-dG-GFP WT and HaCaT cells infected with VSV-dG-RABV-G GFP (MOI=1). **(C).** Graph showing the mean GFP% as quantified by flow cytometry in HaCaT cells infected with VSV-GFP WT and rVSV-dG-RABV-G-GFP. Error bars represented the SEM from three biological replicates (n = 3); ns; non-significant p>0.05, ****, p < 0.0001, by one-way ANOVA. **(D)** Heat map showing the mean RABV-G binding percentage of the rVSV-dG-RABV-G-GFP to HaCaT cells transiently expressing individual or combined P.alecto receptors**. (E). Heat** map showing the mean PFU/mL of rVSV-dG-RABV-G-GFP upon expressing individual and combinatorial P.alecto receptors on HaCaT cells*.

To exclude the possibility that HaCaT cells are resistant to infection with all members of *Rhabdoviridae*. We tested their susceptibility to rVSV-GFP WT, the rVSV-dG-RABV-G-GFP (served as a negative control) at MOI=1.0. Virus replication was first assessed by microscopic observation of GFP, it was observed that 24 hours post infection (hpi), GFP was clearly demonstrated on HaCaT cells infected with rVSV-GFP WT **(Fig 1 A),** unlike mock and HaCaT cells infected with rVSV-dG-RABV-G-GFP which did not show any GFP signal. Results obtained from GFP signal quantification, showed a significant increase in the GFP % of the rVSV-GFP WT infected HaCaT cells compared to the mock and cells infected with rVSV-dG-RABV-GFP **(Figs 1 B-C)** p < 0.0001. These findings clearly indicated the ability of HaCaT cells to support the VSV-GFP WT infection, but not rVSV-dG-RABV-G-GFP.

Next, we sought to determine whether the ectopic expression of any of *P.alecto* RABV receptors individually would render HaCaT cells susceptible to rVSV-dG-RABV-G-GFP infection. The HaCat cells were transfected with plasmids encoding each of the *P.alecto* ORF corresponding to RABV receptors. At 48 hr post transfection, cells were infected with rVSV-dG-RABV-G-GFP at MOI =5. Thirty hrs post infection, we expected to observe GFP, but surprisingly, no GFP was observed. For this purpose, we quantified the binding efficiency of the RABV-G with HaCaT cells expressing different receptors. Results obtained from flow cytometry data, showed significantly higher binding of RABV-G to HaCaT cells expressing ITGB1, mGluR2, nAChR and NCAM (**Fig S 2 A-B)**. While binding of RABV-G in p75 expressing cells, showed no significance difference from empty vector **(Fig. S 2 A-B).** Mock HaCaT cells (un transfected, uninfected) and empty vector control (transfected with empty vector and infected) controls were included. Next, we assessed whether this binding enhanced the released virus particles employing, plaque assay. Interestingly, despite no GFP was observed, significantly higher levels of the released virus progeny were shown in HaCaT cells expressing nAChR, mGluR2 and however, the released virus observed in HaCaT cells ectopically expressing, ITGB1, NCAM and p75**, (Fig S 2 C-D)** showed no significant difference compared to the empty vector control. These results suggested the potential role of ITGB1, nAChR and mGluR2 in promoting RABV entry **(Fig 1 D)**. Given that overexpression of individual receptors in HaCaT cells did not result in GFP observation we further examined whether the simultaneous co-expression of two receptors would render HaCaT cells susceptible to infection by rVSV-dG-RABV-G-GFP.

We introduced two receptors simultaneously into HaCaT cells at equal concentrations. Following a 48-hour post-transfection, cells were infected with rVSV-dG-RABV-G-GFP at an MOI of 5. After 30 hpi, no GFP expression was detected. Consequently, we employed flow cytometry approach to quantify the binding of surface G protein to HaCaT cells expressing the combined receptors. Binding affinity of RABV-G was significantly higher in HaCaT cells ectopically expressing receptor combinations involving mGluR2, ITGB1, nAChR., and NCAM receptors **(Fig 1 D, Fig S 3 A-B)** compared to empty vector, negative control. Non-significant binding capacity was observed on HaCaT cells expressing combination with p75, except when ITGB1 or mGluR2 were combined with p75 they showed significance difference form empty vector control **(Fig S 3 A-B)**.

All mGluR2, ITGB1, NCAM and nAChR receptor combinations showed significantly enhanced viral release compared to EV control **(Fig 1 E, Fig S 4 A-B).** These findings supported that enhanced virus entry and release are promoted when co-expressing receptors simultaneously, with ITGB1 playing a potential role in virus initial attachment. Nevertheless, when combined with p75, the receptor combinations exhibited non-significant binding affinity and released virus compared to the empty vector control **(Fig 1 F, Fig S 4 A-B).** These findings align with previous study that indicated the non-essential role of p75 in RABV entry (8).

To summarize, combinations that included mGluR2, ITGB1, and nAChR receptors exhibited increased binding with the RABV surface glycoprotein. While combinations with the p75 receptor resulted in reduced released viral particles and demonstrated the lowest binding to RABV-G. Additionally, the significant increase in the progeny virus was achieved through combinations involving the mGluR2, nAChR and NCAM. The ectopic expression of nAChR or mGluR2 receptors individually, significantly enhanced the binding capacity with the RABV-G, even exceeding the affinity observed when these receptors were co-expressed. However, none of these receptors were able to render HaCaT cells susceptible to infection as indicated by absence of the GFP signal.

Considering that our study primarily focuses on RABV entry, the selection of ITGB1, mGluR2, and nAChR receptors for further studies was based on their potential role in facilitating RABV-G entry. Further investigation into their specific roles in RABV entry among different species along with knockout studies.

### 3.2 *P.alecto* nAChR receptor enhances virus replication and internalization on Pa-BR cells

Vampire bats have been considered the primary source of RABV transmission in the Americas (25). We tested the possibility of other bat species such as Australian black flying fox (*P.alecto*) for allowing rVSV-dG-RABV-G-GFP replication *in vitro*. Initially, *P.alecto* ORF sequences corresponding to RABV receptors (ITGB1, mGluR2 and nAChR) were retrieved from the NCBI. Sequences were then codon optimized and cloned into pCAGG plasmids with the FLAG tag at the C-terminus with validating their expression and cellular localization by western blot and IFA; respectively **(Fig 2 A - B).** The role of *P.alecto* RABV receptors on rVSV-dG-RABV-G-GFP replication was evaluated during entry and internalization steps. For entry assay, each of *P.alecto* receptor plasmids were ectopically expressed on Pa-BR cells. After 48 hpt, cells were infected with rVSV-dG-RABV-GFP (MOI 5) for 1 hr. Followed by FC analysis with dual labelling infected and transfected cells against rabies G protein and anti-FLAG antibodies; respectively to allow gating the cells bound to RABV-G (GFP+) cells from transfected cells (RFP+). Our FC analysis showed efficient binding of RABV-G to Pa-Br cells over expressing *P.alecto* nAChR and mGluR2 receptors, which resulted in allowing more virus entry. However, cells expressing ITGB1 bat receptor allowed restricted viral entry albeit, significantly different from empty vector control **(Fig 2 C and E).** Further understanding of whether same receptors utilized in RABV entry would allow virus internalization was carried out through quantifying the GFP % after 30 hpi of the transfected cells with rVSV-dG-RABV-G-GFP. Interestingly, our results demonstrated that Pa-Br cells ectopically expressing *P.alecto* nAChR and mGluR2 receptors enhanced the virus replication, compared to the *P.alecto* ITGB1 **(Fig 2 D and F, Fig S 5).**

**Figure 2.**
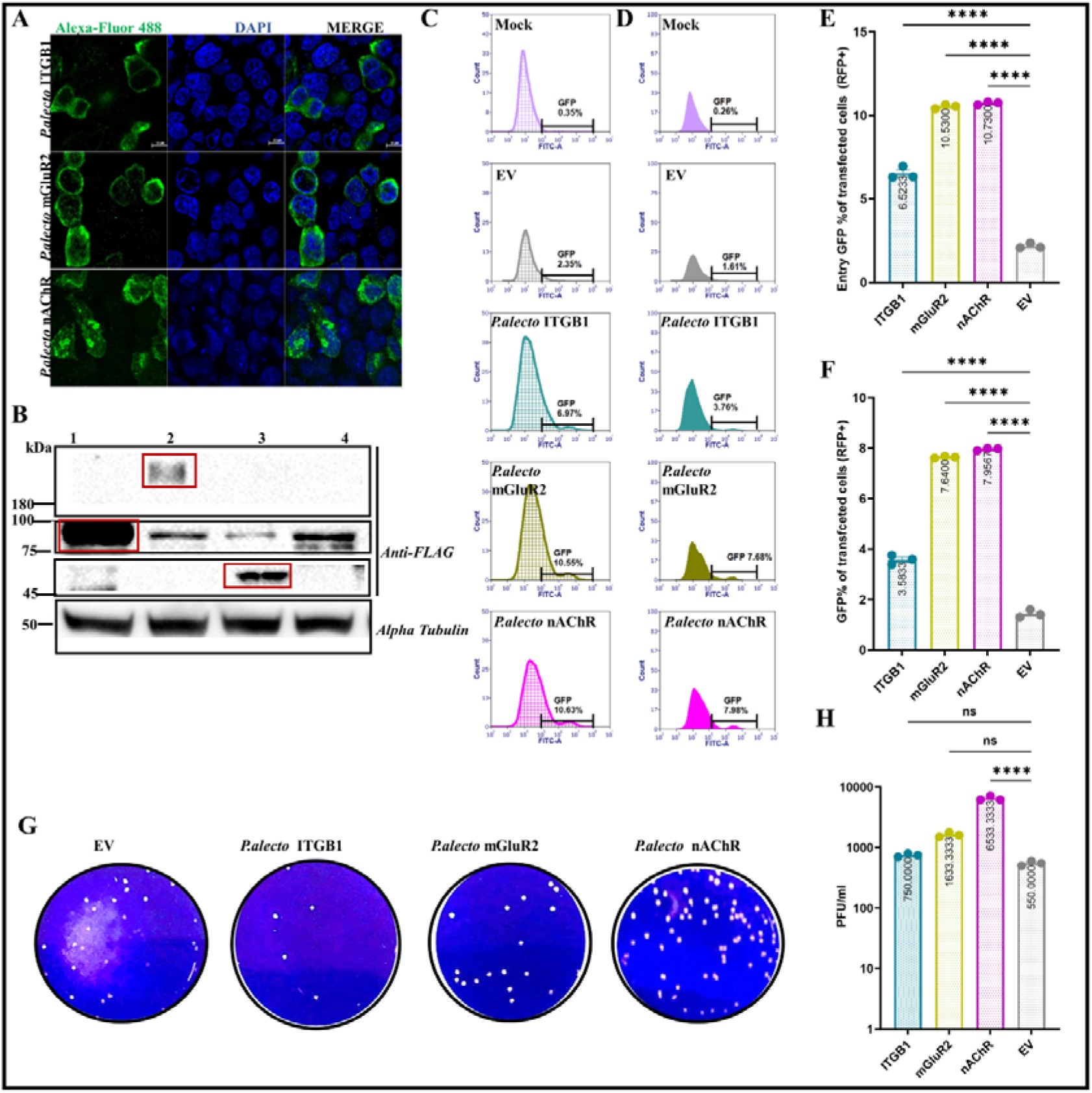
*P.alecto* nAChR enhances entry and replication of rVSV-dG-RABV-G-GFP on Pa-Br cells. **(A)** *The HEK293 cells were transfected with plasmids encoding. P.alecto ITGB1-FLAG, mGluR2-FLAG, and nAChR-FLAG; respectively for 24 h. After 24 hpi, cells were fixed and stained with Anti-FLAG antibody and Alexa-Fluor 468 secondary antibody (Green). Cell nuclei were stained by DAPI (Blue). Fluorescence signals were visualized by confocal immunofluorescence microscopy. Scale bars size, 20* μ*m, images analysed using the ZenCore 3.4 software **(B)** Immunoblot analysis of P.alecto receptors. Pa-Br were transiently transfected with P.alecto receptor plasmids. Thirty-six hrs post-transfection, cell lysates were obtained and subjected to SDS-PAGE and Western blot analysis. The blots were stained against the FLAG tag. Alpha tubulin blot was used as a loading control. The FLAG-tagged cellular proteins showed the expected sizes pCAGG human ITGB1 (expected size: 88*L*kDa), pCAGG human mGluR2 (expected size 190-200 kDa), pCAGG human nAChR (expected size 45*L*kDa) Cell lysates served as negative control. Uncropped blots are shown in Fig S 5 A-B. **(C).** Representative histograms showing the GFP % of rVSV-dG-RABV-G-GFP infected Pa-Br cells. The Pa-Br cells were transiently transfected with the P.alecto receptors and empty vector. Fourty-eight hr post transfection, cells were infected with the rVSV-dG-RABV-G-GFP MOI=5 for 2 hrs., then cells were washed, collected, and stained with anti-FLAG (targeting the FLAG-tagged receptor) and RABV-G antibodies(targeting the virus RABV-G), followed by staining with Alexa Fluor 568 and Alexa Fluor 468 conjugated antibodies, respectively for flow cytometry analysis. Flow cytometry data were analysed by FCS Express software. Cell control represents the un-infected cells. (**D)**. Representative histograms showing the GFP % of rVSV-dG-RABV-G-GFP infected Pa-Br cells. Pa-Br cells were transiently transfected with P.alecto receptors. Forty-eight hrs post transfection, cells were infected with rVSV-dG-RABV-G-GFP, MOI=5. Thirty hpi, the cells collected and stained against the FLAG antibody (targeting the receptors) and followed by staining with Alexa Fluor 568 conjugated antibody for Flow cytometry analysis. GFP% was calculated from the receptor expressing cells, the empty vector transfected Pa-Br cells was used as control, cell control represents the un-infected cell control. **(E).** Graph showing the mean GFP% of Pa-Br cells infected and transfected with P.alecto receptors and EV. The GFP % corresponds to the RABV-G bound to the transfected Pa-Br cells. Data are representative of the mean and SEM of three biological replicates using one way ANOVA. ****, P < 0.0001. **(G)** Graph showing the mean GFP% of Pa-Br cells infected and transfected with P.alecto receptors compared to cells infected and transfected Pa-Br cells with the empty vector. The GFP % corresponds to the rVSV-dG-RABV-G-GFP internalized to the transfected Pa-BR cells. Data are representative of the mean and SEM of three biological replicates using one way ANOVA. ****, P < 0.0001. The experiment was performed three times (n=3) independently. **(H).** Representative plaque morphology of infected Pa-Br cells. Pa-Br cells transiently expressing the P.alecto receptors, were infected with the rVSV-dG-RABV-G-GFP MOI=5. Thirty hpi, the viral supernatants were collected for quantifying the released progeny virus. The released viruses were quantified using plaque assay on BHK-21 cells after 72 hrs. **(H)**. Graph showing the difference of the mean PFU/mL of rVSV-dG-RABV-G-GFP between Pa-Br expressing P.alecto receptors and the Pa-Br cells infected and transfected with the empty vector. Data are representative of the mean and SEM of three biological replicates using one-way ANOVA. ns, non-significant, P > 0.05, ****, P< 0.0001*.

To demonstrate additional evidence of the receptor role in allowing the release of virus progeny, we performed plaque assay. Only Pa-BR cells over expressing *P.alecto* nAChR significantly increased the rVSV-dG-RABV-G-GFP replication compared to empty vector control. While the virus released from Pa-Br cells over expressing *P.alecto* mGluR2 or *P.alecto* ITGB1 receptors showed no significance difference compared to empty vector control **(Fig 2 G-H).**

Taken together, these results support that rVSV-dG-RABV-G-GFP efficiently bind to each of the *P.alecto* nAChR and mGluR2 receptors, allowing enhanced virus entry, replication compared to *P.alecto* ITGB1 receptor.

### 3.3 A549 cells expressing *H.sapiens* ITGB1 enhances viral replication and *H.sapiens* nAChR facilitated virus replication

To test whether *H.sapiens* ITGB1 (NM_002211.4), mGluR2(NM_000839.5) and nAChR (NM_001039523.3) orthologs mediate the rVSV-dG-RABV-G-GFP entry and replication in similar pattern as of *P.alecto* receptors. We codon optimized and cloned each of the full length ORF of *H.Sapiens* ITGB1, mGluR2 and nAChR with FLAG tag at the C-terminus in pCAGG vector. The vectors were referred to as *H.sapiens* ITGB1 (huITGB1), *H.sapiens* mGluR2 (humGluR2) and *H.sapiens* nAChR (hunAChR), respectively. The expression of the plasmids was evaluated by western blot and IFA **(Fig 3 A - B).**

**Figure 3.**
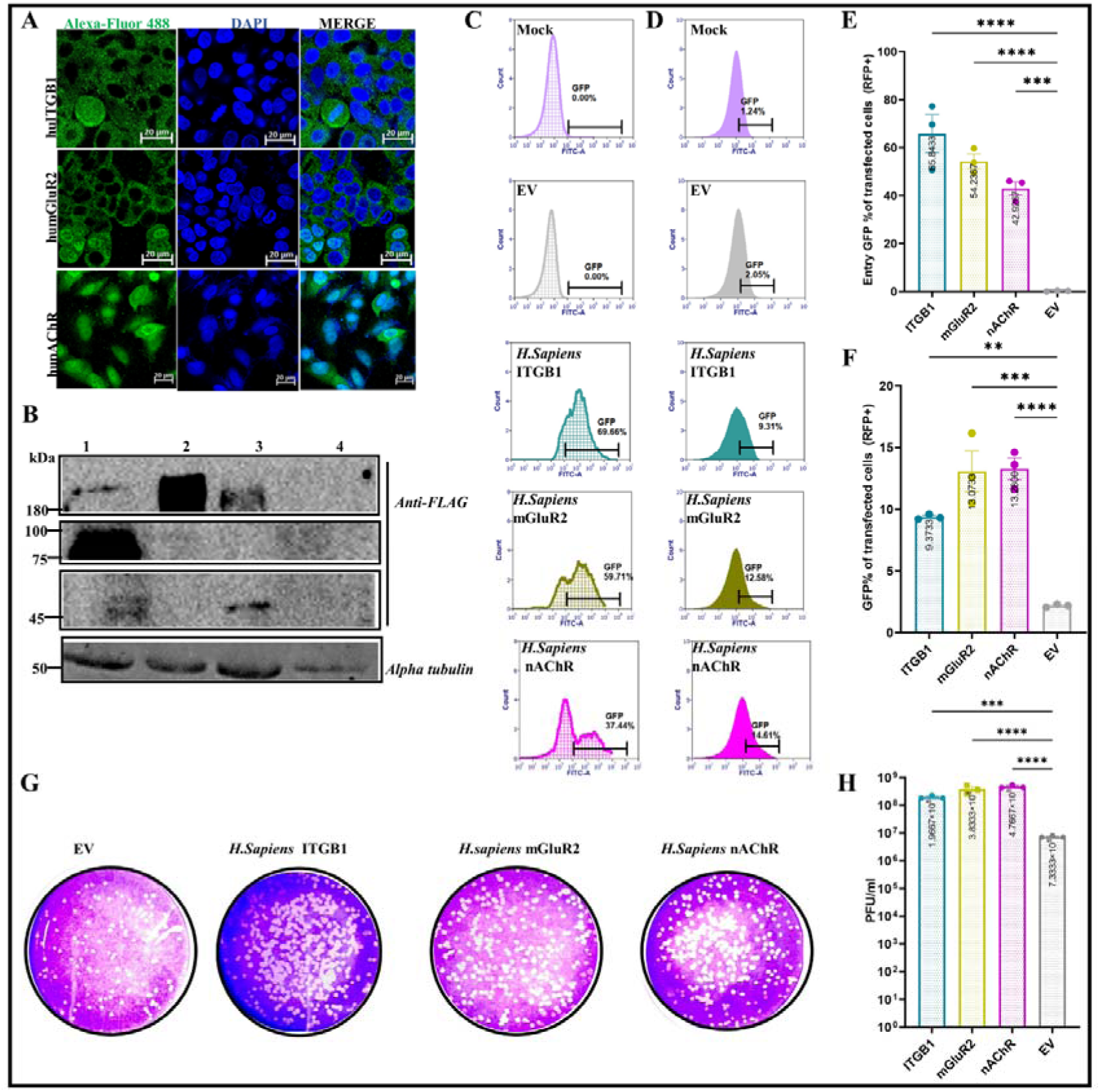
H. sapiens ITGB1 enhances rVSV-dG-RABV-G-GFP entry and nAChR promotes RABV internalization on A549 cells. ***(A)*** *A549 cells were transfected with plasmids encoding. H. Sapiens ITGB1-FLAG, mGluR2-FLAG, and nAChR-FLAG; respectively for 24 h. After 24 hpi, cells were fixed and stained with Anti-FLAG antibody and Alexa-Fluor 468 secondary antibody (Green). Cell nuclei were stained by DAPI (Blue). Fluorescence signals were visualized by confocal immunofluorescence microscopy. Scale bars size, 20* μ*m, images analysed using the ZenCore 3.4 software **(B).** Immunoblot analysis of human receptors. A549 were transiently transfected with H.sapiens receptor plasmids. Thirty-six hrs post-transfection, cell lysates were obtained and subjected to SDS-PAGE and Western blot analysis. The blots were stained against the FLAG tag. Alpha tubulin blot was used as a loading control. The FLAG-tagged cellular proteins showed the expected sizes pCAGG human ITGB1 (expected size: 88*L*kDa), pCAGG human mGluR2 (expected size 190-200 kDa), pCAGG human nAChR (expected size 45*L*kDa) The experiments were performed two times independently (n=2). Cell lysates served as negative control. Uncropped blots are shown in Fig S 6 A-B. **(C).** Representative histograms showing the GFP % of rVSV-dG-RABV-G-GFP infected A549 cells. The A549 cells were transiently transfected with the H.sapiens receptors and empty vector (as control). Forty-eight hr post transfection, cells were infected with the rVSV-dG-RABV-G-GFP MOI=5 for 2 hrs., then cells were washed, collected, and stained with anti-FLAG (targeting the FLAG-tagged receptor) and RABV-G antibodies(targeting the virus RABV-G), followed by staining with Alexa Fluor 568 and Alexa Fluor 468 conjugated antibodies, respectively for flow cytometry analysis. Flow cytometry data were analysed by FCS Express software. Cell control represents the un-infected cells. **(D).** Representative histograms showing the GFP % of rVSV-dG-RABV-G-GFP infected Pa-Br cells. A549 cells were transiently transfected with H.sapiens receptors. Forty-eight hrs post transfection, cells were infected with rVSV-dG-RABV-G-GFP, MOI=5. Thirty hpi, the cells collected and stained against the FLAG antibody (targeting the receptors) and followed by staining with Alexa Fluor 568 conjugated antibody for Flow cytometry analysis. GFP% was calculated from the receptor expressing cells, the empty vector transfected A549 cells was used as control, cell control represents the un-infected cells. **(E).** Graph showing the mean GFP% of A549 cells infected and transfected with H.sapiens receptors and EV. The GFP % corresponds to RABV-G bound to transfected A549 cells. Data are representative of the mean and SEM of three biological replicates using one way ANOVA. **, P < 0.01, ***, P < 0.001. **(F).** Graph showing the mean GFP% of A549 cells infected and transfected with H.sapiens receptors compared to A549 cells infected and transfected with the empty vector. The GFP % corresponds to the rVSV-dG-RABV-G-GFP internalized to the transfected A549 cells. Data are representative of the mean and SEM of three biological replicates using one way ANOVA. **, P < 0.01, ****, P < 0.0001. **(G).** Representative plaque morphology of infected A549 cells. A549 cells transiently expressing the H.sapiens receptors, were infected with the rVSV-dG-RABV-G-GFP MOI=5. Thirty hpi, the viral supernatants were collected for quantifying the released progeny virus. The released viruses were quantified using plaque assay on BHK-21 cells after 72 hrs. **(H).** Graph showing the difference of the mean PFU/mL of rVSV-dG-RABV-G-GFP between A549 cells expressing H.sapiens receptors and the empty vector. Data are representative of the mean and SEM of three biological replicates using one way ANOVA. ***, P < 0.001, ****, P <0.0001*

Next, we assessed the possibility of more efficient use of human RABV receptors by the rVSV-dG-RABV-G-GFP. The A549 cells were transiently transfected with the human vectors encoding each of the RABV receptors (ITGB1, mGluR2 and nAChR) followed by infecting the cells 48 h later with the VSV-dG-RABV-G-GFP (MOI=5.0) for 1 hr, then cells were collected and subjected for FC analysis. Our results demonstrated significant increase in binding affinity of RABV-G to the tested *H.sapiens* receptors compared to EV control, with the highest virus binding observed in cells over expressing the *H.sapiens* ITGB1 **(Fig 3 C and E).**

To identify whether the enhanced RABV-G binding to A549 cells ectopically expressing *H.sapiens* ITGB1 facilitated virus replication, we employed the infection assay. Our results demonstrated significantly higher GFP percentage on cells expressing *H.sapiens* nAChR and mGluR2 compared to cells transfected with *H.sapiens* ITGB1. The obtained results showed significant difference from the EV control **(Fig 3 D and E, Fig S 6)**. These findings were further supported by quantifying the released virus from cells using plaque assay. Of the three tested receptors, all infected cells resulted in significantly higher release of the progeny virus compared to the empty vector, with the highest virus progeny release on human cells transfected with the *H.sapiens* nAChR and mGluR2 **(Fig 3 G and H).**

### 3.4 ITGB1 enhances virus attachment and replication on MDCK cells

To test RABV receptor preference on MDCK cells, the full length ORF of *C.familiaris* ITGB1 (XM_038658027.1), mGluR2 (XM_038427610.1) and nAChR (NM_001003144.2) were codon optimized, synthesized, and cloned in pCAGG vector with a FLAG tag at the C-terminus, referred to as *C.familiaris* ITGB1 (doITGB1), *C.familiaris* mGluR2 (domGluR2) and *C.familiaris* nAChR (donAChR). The localization of the canine receptors and expression were assessed **(Fig 4 A-B)**.

**Figure 4.**
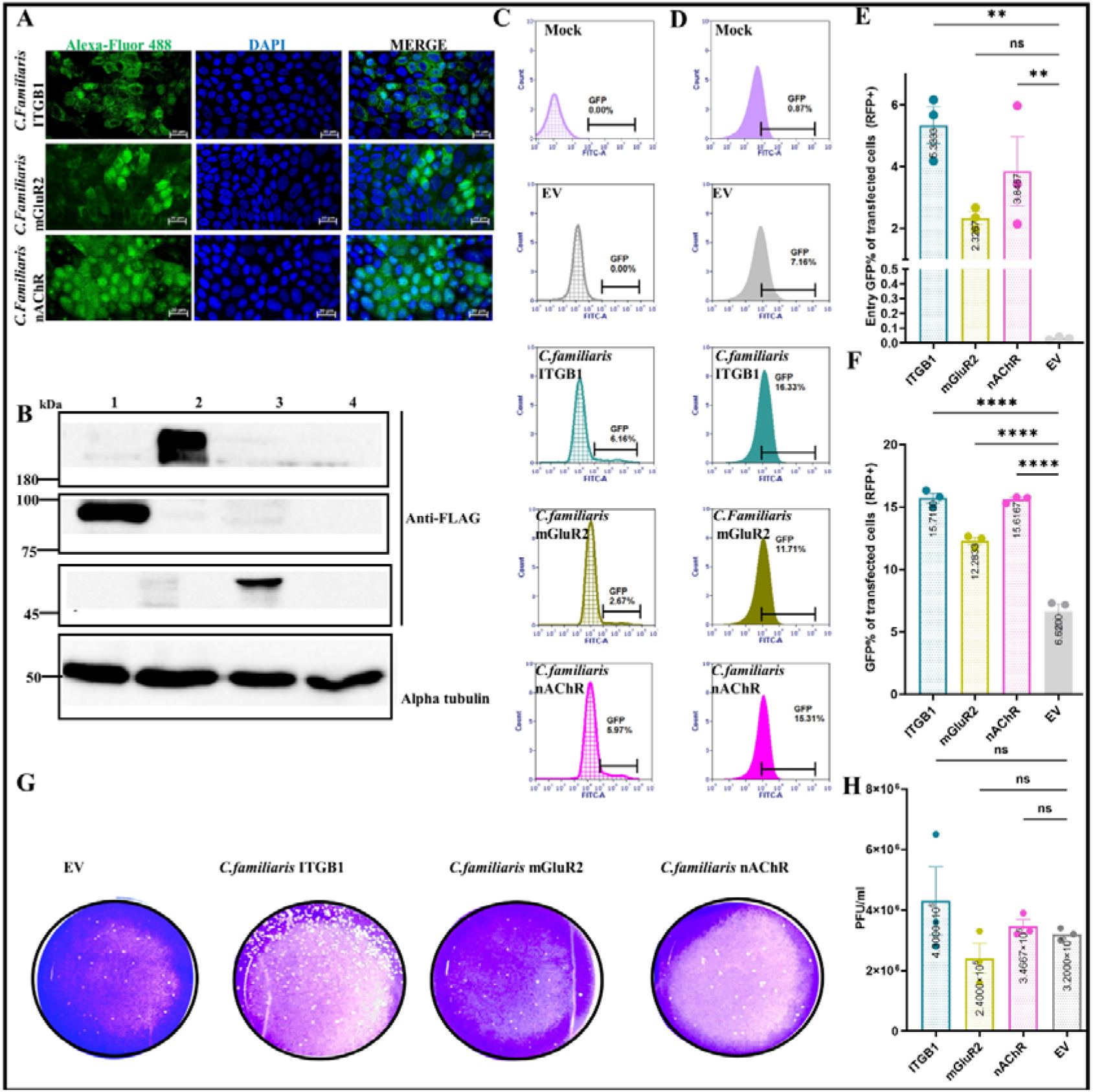
C. familiaris ITGB1 promoted rVSV-dG-RABV-G-GFP entry and replication on MDCK cells. ***(A)*** *MDCK cells were transfected with plasmids encoding. C.familiaris ITGB1-FLAG, mGluR2-FLAG, and nAchR-FLAG; respectively for 24 h. After 24 hpi, cells were fixed and stained with Anti-FLAG antibody and Alexa-Fluor 468 secondary antibody (Green). Cell nuclei were stained by DAPI (Blue). fluorescence signals were visualized by confocal immunofluorescence microscopy. Scale bars size, 20* μ*m Images analysed using the ZenCore 3.4 software. **(B).** Immunoblot analysis of dog receptors on MDCK cells. MDCK cells were transiently transfected with the C.familiaris receptor plasmids. Thirty-six hrs post-transfection, cell lysates were obtained and subjected to SDS-PAGE and Western blot analysis. The blots were stained against the FLAG tag. Alpha tubulin blot was used as a loading control. The FLAG-tagged cellular proteins showed the expected sizes pCAGG canine ITGB1 (expected size: 88*L*kDa), pCAGG canine mGluR2 (expected size 180-200 kDa), pCAGG canine nAChR (expected size 45*L*kDa). Cell lysates served as the negative control. The experiment was performed two times independently (n=2)*. *Uncropped blots are shown in Fig S 7 A-C. **(C)**. Representative histograms showing the GFP % of rVSV-dG-RABV-G-GFP infected MDCK cells. The MDCK cells were transiently transfected with the C.familiaris receptors and empty vector. Fourty-eight hr post transfection, cells were infected with the rVSV-dG-RABV-G-GFP MOI=5 for 2 hrs., then cells were washed, collected, and stained with anti-FLAG (targeting the FLAG-tagged receptor) and RABV-G antibodies(targeting the virus RABV-G), followed by staining with Alexa Fluor 568 and Alexa Fluor 468 conjugated antibodies, respectively for flow cytometry analysis. Flow cytometry data were analysed by FCS Express software. Cell control represented the un-infected cells. (**D).** Representative histograms showing the GFP % of rVSV-dG-RABV-G-GFP infected MDCK cells. MDCK cells were transiently transfected with C.familiaris receptors. Forty-eight hrs post transfection, cells were infected with rVSV-dG-RABV-G-GFP, MOI=5. Thirty hpi, the cells collected and stained against the FLAG antibody (targeting the receptors) and followed by staining with Alexa Fluor 568 conjugated antibody for Flow cytometry analysis. GFP% was calculated from the receptor expressing cells, the empty vector transfected MDCK cells was used as control, cell control represented the un-infected cells. **(E).** Graph showing the mean GFP% of MDCK cells infected and transfected with C.familiaris receptors and EV. The GFP % corresponds to the RABV-G bound to the transfected MDCK cells. Data are representative of the mean and SEM of three biological replicates using one way ANOVA. ****, P* ≤ *0.0001**. (F)**. Graph showing the mean GFP% of MDCK cells infected and transfected with C.familiaris receptors compared to cells infected and transfected with the empty vector. The GFP % corresponds to the rVSV-dG-RABV-G-GFP internalized to the transfected MDCK cells. Data are representative of the mean and SEM of three biological replicates using one way ANOVA. *, P* ≤ *0.05, ***, P* ≤ *0.001, ****, P* ≤ *0.0001. **(G).** Representative plaque morphology of infected MDCK cells. MDCK cells transiently expressing the C.familiaris receptors, were infected with rVSV-dG-RABV-G-GFP MOI=5. Thirty hpi, the viral supernatants were collected for quantifying the released progeny virus. The released viruses were quantified using plaque assay on BHK-21 cells after 72 hrs. **(H).** Graph showing the difference of the mean PFU/mL of rVSV-dG-RABV-G-GFP between MDCK expressing C.familiaris receptors and the MDCK cells infected and transfected with the empty vector. Data are representative of the mean and SEM of three biological replicates using one way ANOVA. ns, non-significant, P > 0.05*.

Since dogs represent the main RABV reservoir, we tested the role of canine receptors in rVSV-dG-RABV-G-GFP entry and replication on MDCK cells. To determine the binding preference of the RABV-G to cells expressing canine receptors, the entry assay was employed which showed a substantial increase in viral entry on MDCK cells transiently expressing *C.familiaris* ITGB1, whilst less binding capacity of RABV-G was demonstrated on cells ectopically expressing the *C.familiaris* nAChR. Notably, the lowest RABV-G binding was demonstrated on MDCK cells overexpressing *C.familiaris* mGluR2 which showed no significant difference from EV control **(Fig 4 C and E)**. To evaluate whether RABV employs similar receptor preference during internalization, the GFP percentage and released virus progeny were measured upon rVSV-dG-RABV-G-GFP infection on MDCK cells. Consistent with obtained results from the binding assay, a significant increase in the GFP percentage was observed in cells transiently expressing *C.familiaris* ITGB1, followed by *C.familiaris* nAChR and *C.familiaris* mGluR2 compared to the empty vector control **(Fig 4 D and F, Fig S 7).** The plaque assay results demonstrated a similar trend to FC data, despite showing no significant difference from the EV control **(Fig 4 G - H)**.

Collectively, the obtained results indicated the significant role of *C.familiaris* ITGB1 in mediating the entry and replication of rVSV-dG-RABV-G-GFP on MDCK cells.

### 3.5 Impaired rVSV-dG-RABV-G-GFP entry on ITGBI KO cells

Given the potential role demonstrated by ITGB1 and mGluR2 in rVSV-dG-RABV-G-GFP entry and replication demonstrated from receptor preference experiments on HaCaT cells. We attempted to knockout ITGB1 and mGluR2 to define their exact role in RABV entry and replication.

ITGB1 has been recently identified to facilitate RABV peripheral entry (1). To investigate the function of ITGB1 during RABV replication, we adopted the CRISPR/Cas9 mediated knockout in which a sgRNA targeting the fourth exon of ITGB1 gene was and cloned in PX459 V2.0 vector **(Fig S 9)**. Upon selection and expansion of the single cell colonies, the genomic DNA of the single cell KO clone and WT cells were extracted and the region encompassing the sgRNA was amplified by PCR for band size comparison and sequence analysis. A smaller band corresponding to the KO single cell clone was observed compared to the WT. Further PCR sequencing of the ITGB1 single cell clones originated from KO A549 cells analysis showed the derivation of two CRISPR indels in the KO single cell clones. The deletion generated two different alleles with deletion of either three bases or twenty bases induced in the targeted fourth exon of ITGB1 **(Fig S 9)**. Functional studies were carried out to further elucidate the potential impact of ITGB1 absence in rVSV-dG-RABV-G-GFP replication. The ITGB1 KO cell, ITGB1 KO cells over expressing the human ITGB1 cDNA together with the WT A549 cells were infected with rVSV-dG-RABV-G-GFP (MOI 5) for 24 hr, a KO uninfected cell control was employed. Next, we evaluated the rVSV-dG-RABV-G-GFP infectivity on the cells through visualizing and measuring the GFP percentage. A significant reduction in the GFP percentage was shown on infected ITGB1 KO cells with EV (non-significantly different from the uninfected KO control) compared to WT cells. Over expressing the human ITGB1 on ITGB1 KO cells showed increase in GFP levels, however, the GFP expression levels remained non significantly different from infected ITGB1 KO cells **(Fig 5 A-B, Fig S 9)**. Quantification of released virus progeny showed significant reduction of released virus progeny on ITGB1 KO cells by 3-fold, which was significantly less compared to WT cells. However, the over expression of huITGB1 was capable of recapitulating the rVSV-dG-RABV-G-GFP replication as in the WT cells **(Fig 5 C-D)**. To test whether the efficiency of RABV-G entry differs in the absence and presence of ITGB1, we infected ITGB1 KO and ITGB1 KO cells expressing huITGB1 with rVSV-dG-RABV-G-GFP MOI=5 for 1 hr to allow attachment of RABV-G to susceptible cells, followed by evaluating the binding percentage. The RABV-G binding to KO ITGB1 cells showed no significant difference from the KO uninfected cell control. However, significantly enhanced virus binding was detected in ITGB1 KO cells transiently expressing the human ITGB1 receptor **(Fig 5 E-F)**. These results suggest the potential role of ITGB1 in promoting RABV entry.

**Figure 5.**
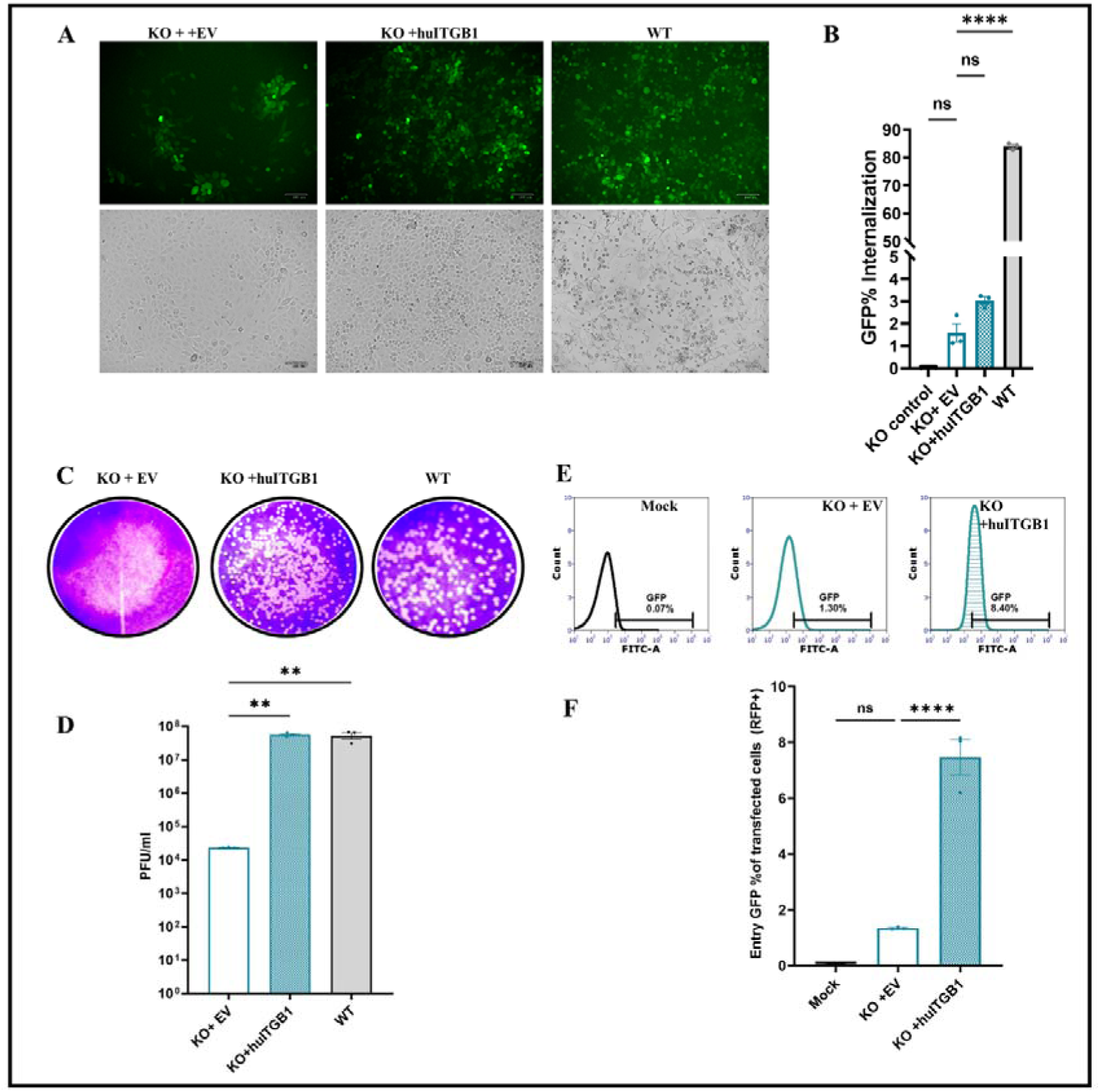
Impaired rVSV-dG-RABV-G-GFP entry on ITGB1 KO cells. ***(A).*** *Representative microscopic fields green (left), bright (right) for A549 KO ITGB1 cells infected with rVSV-dG-RABV-G-GFP MOI=5. A549 KO ITGB1 cells, were transfected with empty vector and human ITGB1 FLAG vector; respectively, after 48h, cells were infected with rVSV-dG-RABV-G-GFP (MOI=5). Thirty hpi, the GFP percentage corresponding to virus replication were imaged. The WT A549 cells were used as infection control **(B)**. Graph showing the mean GFP% of ITGB1 KO cells transiently transfected with empty vector and human ITGB1 FLAG vectors; respectively. The KO control cells were used as non-infected control and the WT A549 cells were used as infection control. The GFP% corresponds to the internalized virus. Error bars represent the SEM of three biological replicate samples. using one-way ANOVA. ns, non-significant, P > 0.05., **** P* ≤ *0.0001. **(C)**. Representative plaque morphology of A549 KO ITGB1 cells infected with rVSV-dG-RABV-G-GFP. The A549 KO ITGB1 cells were transiently transfected with the empty vector and human ITGB1 FLAG, respectively. Forty-eight hr post-transfection, cells were infected with the rVSV-dG-RABV-G-GFP MOI=5. Thirty hpi, the viral supernatants were collected for quantifying the released progeny virus. The released viruses were quantified using plaque assay on BHK-21 cells after 72 hrs. **(D).** Graph showing the difference of the mean PFU/mL of rVSV-dG-RABV-G-GFP between A549 ITGB1 KO cells transfected with either EV or human ITGB1 FLAG, the WT cells were used as infection control and KO cell used as non-infected control. **(E).** Representative histograms showing the GFP % of rVSV-dG-RABV-G-GFP infected A549 KO ITGB1 cells. The A549 KO ITGB1 cells were transiently transfected with the empty vector and human ITGB1 FLAG, respectively. Forty-eight hr post-transfection, cells were infected with the rVSV-dG-RABV-G-GFP MOI=5 for 2 hrs. Then cells were collected and stained with anti-FLAG (targeting the FLAG-tagged receptor) and RABV-G antibodies (targeting the virus RABV-G), followed by staining with Alexa Fluor 568 and Alexa-Fluor 468 conjugated antibodies, respectively for flow cytometry analysis. Flow cytometry data were analysed by FCS Express software. **(F)**. Graph showing the mean GFP% of A549 KO ITGB1 cells infected and transfected with EV and human ITGB1 vector, KO control was used as non-infected control The GFP % corresponds to the RABV-G bound to the transfected A549 KO ITGB1 cells. Data represent the average of three biological replicates with S.E.M. using one-way ANOVA. ns, non-significant, P > 0.05., ** P* ≤ *0.01, **** P* ≤ *0.0001*

### 3.6 A549 KO mGluR2 significantly reduced rVSV-dG-RABV-G-GFP released virus progeny

To investigate whether the complete deletion of mGluR2 gene would still allow the VSV-dG-RABV-G-GFP replication in A549 cells. We adopted CRISPR/Cas9 mediated knockout strategy in which single guide RNA (sgRNA) targeting the second exon of the mGluR2 was designed and cloned in PX459 V2.0 vector for simultaneous expression of the Cas9 endonuclease and the sgRNA upon transfection in the in A549 cells. Analysis of the generated mGluR2 KO and WT A549 genomic DNA from cells clones was carried out initially by amplifying the genomic sequence flanking 200 bases upstream and 200 bases downstream the designed sgRNA which revealed a difference in size of KO clone compared to the WT cell clones **(Fig S 10)**. Owing to the unavailability of the commercial antibodies specific to the mGluR2 protein in our lab, validation of the KO cells was carried out through sequencing and functional analysis. Sequence analysis of both the PCR products from KO and WT cell clones sequence showed a variety of indels in mGluR2 exon 2. The resulted indels included clones with deleted nucleotides and other clones showed inserted nucleotides compared to the WT sequence **(Fig S 10).**

To determine to which extent mGluR2 KO cells would support RABV replication, the following cells were infected with rVSV-dG-RABV-GFP at MOI of 5: mGluR2 KO cells over expressing the human mGluR2, mGluR2 KO cells overexpressing empty vector and WT A549 cells, a KO uninfected cell control was employed. After 30 hpi, GFP was observed RABV in both KO and WT cells, however less signal was clearly observed in mGluR2 KO cells **(Fig 6 A).** The percentage of internalized virus was measured by FC analysis which showed significantly lower GFP percentage in mGluR2 KO cells compared to WT cells. Ectopic expression of human mGluR2 on KO mGluR2 allowed significantly higher levels of the internalized virus compared to mGluR2 KO infected cells **(Fig 6 B, Fig S 11)**. Comparing the levels of the released virus particles, indicated significant inhibition of the rVSV-dG-RABV-G-GFP replication in mGluR2 KO cells by 5-fold compared to WT cells. Interestingly, the expression of humGluR2 in KO cells was capable of restoring the viral replication as the WT cells which was significantly elevated compared to mGluR2 KO cells **(Fig 6 C-D)**. To dissect at which stage of RABV replication the mGluR2 is required, an entry assay was performed to compare the entry of rVSV-dG-RABV-G-GFP on mGluR2 KO cells and mGluR2 KO cells ectopically expressing the human mGluR2. FC analysis for evaluating the RABV-G entry showed that mGluR2 KO cells allowed significantly higher initial binding of the RABV-G compared to uninfected cell control, yet it was significantly less compared to mGluR2 KO cells expressing humGluR2 **(Fig 6. E-F)**. Collectively, the depletion of mGluR2 levels on A549 cells, resulted in significant reduction of RABV replication and reduced RABV-G capacity of to bind to the cells. These findings suggest the potential role of the mGluR2 in rVSV-dG-RABV-G-GFP entry and replication, yet its absence did not completely inhibit virus replication.

**Figure 6.**
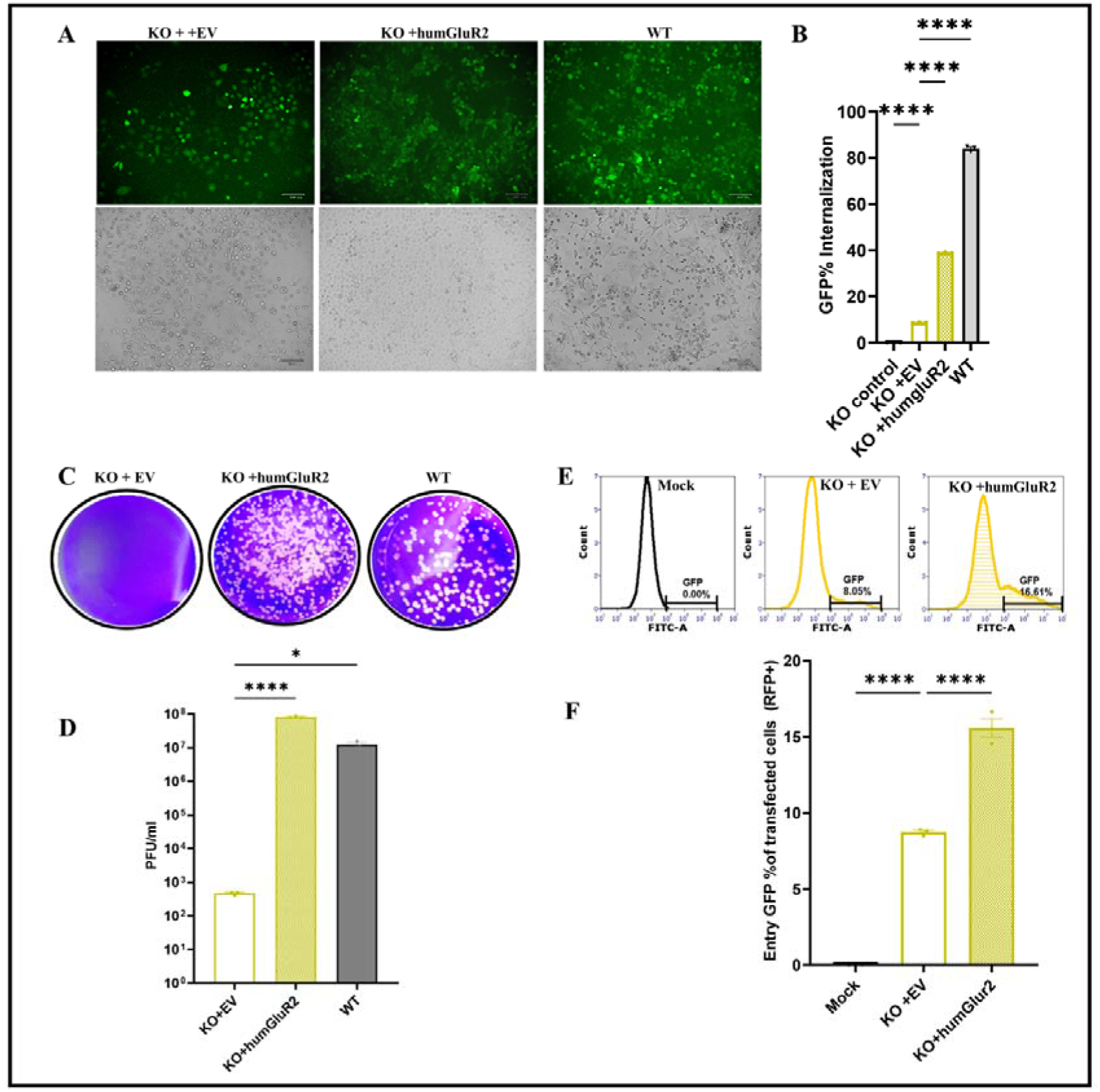
Reduced replication of rVSV-dG-RABV-G-GFP on mGluR2 KO cells. ***(A). Representative*** *microscopic fields green (left), bright (right) for A549 KO mGluR2 cells infected with rVSV-dG-RABV-G-GFP MOI=5. A549 KO mGluR2 cells, were transfected with empty vector and human mGluR2 FLAG vector; respectively, after 48h, cells were infected with rVSV-dG-RABV-G-GFP (MOI=5). Thirty hpi, the GFP percentage corresponding to virus replication were imaged, The WT A549 cells were used as infection control **(B).** Graph showing the mean GFP% of mGluR2 KO cells transiently transfected with empty vector and human mGluR2 FLAG vectors; respectively. The KO control cells were used as, non-infected control and the WT A549 cells were used as infection control. The GFP% corresponds to the internalized virus. All experiments were performed three times (n=3) independently. Error bars represent the SEM of three biological replicate samples. using one-way ANOVA. **** P* ≤ *0.0001. **(C).** Representative plaque morphology of the A549 KO mGluR2 cells infected with rVSV-dG-RABV-G-GFP. The A549 KO mGluR2 cells were transiently transfected with the empty vector and human mGluR2 FLAG, respectively. Forty-eight hr post-transfection, cells were infected with the rVSV-dG-RABV-G-GFP MOI=5. Thirty hpi, the viral supernatants were collected for quantifying the released progeny virus. The released viruses were quantified using plaque assay on BHK-21 cells after 72 hrs**. (D).** Graph showing the difference of the mean PFU/mL of rVSV-dG-RABV-G-GFP between A549 mGluR2 KO cells transfected with either EV or human mGluR2 FLAG, the WT A549 cells were used as infection control. **(E).** Representative histograms showing the GFP % of rVSV-dG-RABV-G-GFP infected A549 KO mGluR2 cells. The A549 KO mGluR2 cells were transiently transfected with the empty vector and human mGluR2 FLAG; respectively. Forty-eight hr post-transfection, cells were infected with the rVSV-dG-RABV-G-GFP MOI=5 for 2 hrs. After washing with PBS, for removal of unbound virus, cells were collected and stained with anti-FLAG (targeting the FLAG-tagged receptor) and RABV-G antibodies (targeting the virus RABV-G), followed by staining with Alexa Fluor 568 and Alexa Fluor 468 conjugated antibodies, respectively for flow cytometry analysis. Flow cytometry data were analysed by FCS Express software. **(F)**. Graph showing the mean GFP% of A549 KO mGluR2 cells infected and transfected with EV and human mGluR2 vector, KO control was used as non-infected control The entry GFP % corresponds to the RABV-G bound to the transfected A549 KO mGluR2 cells. Data represent the average of three biological replicates with S.E.M. using one-way ANOVA. *, P* ≤ *0.05, ****, P* ≤ *0.0001*.

## 4. Discussion

Entry of RABV into host cells is typically mediated by the interactions of cellular receptors with the surface glycoprotein (6). With the advancement in high throughput technology and genetic screening methods, new proteins have been identified as RABV receptor candidates (1,7,9,10,26). However, this knowledge represents an enormous potential for designing new structural guided antiviral drugs. More understanding of the underlying mechanism by which RABV binds and internalizes to the cellular compartments requires further investigation, besides identification of the major receptor directing RABV entry.

Initially, our aim was to experimentally establish a cell line that is resistant to RABV entry, to dissect the specificity of RABV receptors in mediating its entry. Thus, we infected various cell lines with rVSV-dG-RABV-GFP which showed the wide tropism of rVSV-dG-RABV-G-GFP, exemplified in its ability to replicate in the wide spectrum of tested cell lines. However, a distinct alteration in cell tropism was observed among the VSV-GFP WT and the rVSV-dG-RABV-G-GFP infection on HaCaT cells which revealed resistance to the recombinant virus displaying the RABV-G protein, in contrast to their permissiveness to the rVSV -GFP-WT, suggesting the suitability of HaCaT cells to act as a cellular model for studying RABV receptor preference (27). In our study, we ectopically expressed the *P.alecto* receptors on HaCaT cells (Human cell line) to establish a heterologous system in which other factors that could affect the cellular receptors interactions are eliminated Additionally, to alleviate the cellular stress induced by transfection, we incorporated an empty vector control throughout the entire study as a means of normalization.

The obtained results from individual and combined ectopic expression of the RABV receptors on HaCaT cells showed neither distinguished cytopathic effect (CPE) nor any GFP upon infection with the rVSV-dG-RABV-G-GFP. Thus, comparing the effect of receptors on the cell’s permissiveness was based on the binding percentage with RABV-G protein and the quantification of virus released progeny in the supernatants. Clearly more binding and production of virus progeny was observed on infected HaCaT cells individually expressing nAChR or mGluR2 receptors. Further, the effect of co-expressing receptors with each of mGluR2, nAChR, ITGB1 or NCAM resulted in more enhanced viral replication than combinations involving the p75 receptor. This might be correlated with findings from previous study which showed the possible transport of the RABV in p75 in-dependent pathway. Highlighting that the role of p75 receptor is limited to accelerating RABV transport through internalization (26).

Interestingly, the higher levels of released virus progeny demonstrated on HaCaT cells co expressing the ITGB1 with nAChR receptors is supported by findings from previous study which demonstrated the interaction of both receptors through CO-IP assay (1). Notably, higher levels of virus release were detected in infected HaCaT cells expressing NCAM in combination with mGluR2 or ITGB1 or nAChR. This could be delineated to the role of NCAM in conferring resistant cells susceptible to RABV infection (9).

From HaCaT cell receptor preference studies, several conclusions can be drawn. Firstly, is that cells’ susceptibility to virus infection is not solely governed by the expression of entry receptors. Hence the ectopic expression of RABV receptors on HaCaT cells showed enhanced virus binding and virus release but could not render them susceptible to infection (no GFP was observed). This could be explained by the fact that accumulation or high binding between the surface glycoprotein and the receptor does not necessarily trigger post-binding events that allow viral entry, due to the possible presence of restriction factors or lack of other genes expression required for viral infection (27,28). Another probable reason for the absence of CPE in HaCaT cells infected with rVSV-dG-RABV-GFP might be due to the conservation of HaCaT cell integrity as previously observed in HaCaT cells infected with rhinovirus (29). The receptor preference results on HaCaT cells suggested enhanced binding and replication of RABV upon interacting with nAChR, mGluR2 and ITGB1 receptors. Despite the prominent levels of virus replication on NCAM expressing cells, we did not attempt to further study and genetically delete the NCAM gene. Since it has been previously reported that NCAM deficient mice only exhibited delay in rabies associated mortality and restricted RABV invasion (9,30).

Over the past years, the origin of the Ebola virus, SARS-CoV, MERS-CoV, Hendra virus and other viral diseases have been linked to the bats (31). With specific reference to the bat species belonging to genus Pteropus which have been previously reported as Hendra virus reservoirs (32). Thus, we investigated *P.alecto* brain cell line susceptibility to RABV infection. Remarkably, to the best of our knowledge our study is the first to report that *P.alecto* RABV encoding receptor genes are functional and that Pa-BR cell line which is derived from the *P.alecto* brain could be infected with RABV. This finding is crucial in understanding the potential role that could be played by the *P.alecto* as potential virus reservoir of the RABV wildlife. However, it is worth mentioning that *in vitro* susceptibility of the bat species to the virus does not inevitably imply its susceptibility to the RABV *in vivo* (32). Considering the numerous factors which influence the likelihood of the virus transmission from bats to human in the field, including the bat immune response and the ecological interaction which would allow the contact between the virus and the host and behavioural patterns (31). Thus further *in vivo* studies to understand the significance of *P.alecto* susceptibility to virus infection and the explanation of the restricted RABV reservoirs to the microbat species are deemed essential.

Since most of human rabies cases have originated from cross species transmission events through bats or dogs confirming that human health is connected to animal health, suggesting that one health approach is among the best strategies to control rabies (33). It is worth noting that host cellular factors are essential prerequisites for establishment of infection into a new host (34). Thereby, understanding RABV-G ability to escape the receptor binding specificity among human, bat and canine cell lines employing ITGB1, mGluR2 and nAChR orthologs will contribute to understanding rabies viral transmission (35,36). We achieved this through comparatively testing the role of ectopically expressed RABV receptor orthologs in promoting rVSV-dG-RABV-G-GFP (belongs to dog related phylogeny) ability to enter and replicate *in vitro* along with mapping its receptor preference in each of the selected cell lines in comparison to dog receptors.

Unexpectedly, our results indicated that RABV entry and replication on different cell lines exhibited differential receptor preference. The replication and entry of rVSV-dG-RABV-G-GFP on Pa-Br cells was enhanced by the overexpression of the *P.alecto* nAChR receptor. While on MDCK cells, the virus initial attachment and virus GFP expression were supported by ectopic expression of *C.familiaris* ITGB1. In human cell line, the initial attachment was augmented by *H.sapiens* ITGB1, while the virus replication and release were enhanced by expression of *H.sapiens* nAChR. There are several possible explanations for rVSV-dG-RABV-GFP differential receptor preference among different cell lines. One important observation is that nAChR over expressing human and bat cells resulted in more efficient virus release. This might be attributed to the conserved interaction site of nAChR among those species (37). Interestingly, *P.alecto* ITGB1 expressing Pa-Br cells, showed the lowest levels of viral titres and binding compared to enhanced entry and replication on MDCK cells ectopically expressing *C.familiaris* ITGB1. This might be correlated to differences in critical amino acid as *P.alecto* ITGB1 sequence lacks the integrin plexin domain as previously identified (37,38). One other possible scenario for differential receptor utilization of RABV might be the cascade of events activated following the initial attachment to a specific receptor. For instance, when a virus interacts with surface cellular receptor, it could trigger a signal transduction process, which may lead to secretion of interferons and ultimately hinder the virus from being taken up effectively (39). This phenomenon might elucidate why A549 cells that overexpress ITGB1 showed increased viral binding, while enhanced virus replication was demonstrated on the cells transiently expressing nAChR.

These findings could imply that for the virus to successfully cross into and infect other species, it may undergo evolutionary changes, such as mutations, to enhance its adaptation for receptor binding and ultimately overcoming the species barriers through which the virus expand its host range and ensure its survival (40). It is noteworthy that RABV-G strain employed in this study was associated with dog-related genetic lineage (41). Therefore, future research should account for potential variations elicited from interaction of G protein from a bat-related genetic lineage with receptor orthologs of different species. Taken together, the findings obtained from differential receptor preference among species demonstrated by rVSV-dG-RABV-G-GFP might explain the role of these receptors in enhancing virus release which might be the reason for the virus transmission (36).

Further, we attempted to disrupt the gene composition of each of the ITGB1, mGluR2 and nAChR to elucidate at which stage of RABV replication these receptors were required.

Disrupting the gene is initiated through delivery of sgRNA which is complementary to the target site in gene of interest which when coupled with the Cas9 resulting in gene disruption (42). To this aim we used the sgRNA approach targeting specific coding region in ITGB1, mGluR2 and nAChR genes individually. The genetic characterization of the clonal knockouts of the ITGB1 and mGluR2 cell lines was carried out, which demonstrated multiple alleles upon disrupting the ITGB1 or mGluR2 which showed either deletion or insertion in the targeted exon regions and causing frameshift mutations. Owing to the unavailability of commercial antibodies in our laboratory for the ITGB1 and mGluR2 receptors, further confirmation of the disrupted genes effect on RABV replication was carried out through functional studies.

The results obtained from entry and replication of rVSV-dG-RABV-G-GFP on ITGB1 KO, showed that deficient levels of ITGB1 in A549 cells resulted in significant reduction of G protein binding to cells (entry), despite displaying higher levels of RABV released progeny compared to knockout mGluR2 cell line, suggesting its significant role in the initial cellular attachment to the RABV-G. These results are contradicting the results from a recent study which suggested that ITGB1 is co localized with the RABV in early and late endosomes, indicating its role in virus internalization (1). Thus, more studies are required to elucidate in which step of virus replication the ITGB1 is involved.

In contrast, mGluR2 KO cells demonstrated that rVSV-dG-RABV-GFP replication was more influenced than the virus initial binding. In line with these findings, a previous study showed that mGluR2 is internalized through similar endocytic pathway as that of RABV in which they were shown to be co-internalized into the early and late endosomes, highlighting the role of the mGluR2 in RABV internalization (7). Besides, a previous report which demonstrated survival of 58% mice with mGluR2 knockout against RABV challenge (7). This might be concluded that however KO mGluR2 cells supported initial virus binding, virus replication was substantially inhibited, highlighting the mGluR2 prevalent role in virus internalization. Our attempt to generate nAChR KO cells was not successful.

Overall, it is important to note that virus-receptor interaction represent one aspect of the complex interactions between the virus and host and many other factors which influence the successful establishment of infection into new host species. Among those are the specific virus and host cell line being studied, the expression levels of other cellular factors, and the viral entry pathways which ultimately contribute to the fate of RABV infections.(43).

## 5. Conclusion

In conclusion, our study points out several conclusions, firstly, is that it is tempting to speculate that mGluR2 plays the most crucial role in RABV infection since its depletion from cells resulted in the least virus replication, evidenced by less released virus progeny compared to KO ITGB1 cells. Secondly, RABV internalization might be regulated by interactions between two receptors as has been also reported by a recent study which showed that RABV-mGluR2 complex internalization is dependent on interaction with the transferrin receptor protein enabling the uptake of the RABV-mGluR2 complex in the clathrin coated pits (10). In addition to another study which outlined the potential role of an interaction between ITGB1 and nAChR receptors in enhancing the peripheral entry of RABV (1). Since these results indicated the independence of RABV on the previously identified receptors suggesting that it utilizes these receptors simultaneously rather than sequentially as the knockout of these receptors failed to abolish virus entry/replication (ITGB1, mGluR2). Implying that antiviral drugs targeting RABV receptors would not confer proper virus control as those targeting the virus G protein.

## Funding

This study was funded by the Biotechnology and Biological Sciences Research Council (BBSRC) (BB/M008681/1 and BBS/E/I/00001852) and the British Council (172710323 and 332228521). The Ph.D. studies of MEK have been financially supported by Newton Mosharafa-Fund (Bureau ID: NMM11/19) and the Egyptian Ministry of Higher Education and Scientific Research, Cultural Affairs and Mission Sector, Egypt

## Author Contributions

Conceptualization, M.E.K and M.M.; Formal analysis, M.E.K; Investigation and methodology, MEK; Validation of results and project administration M.E.K and M.M; Resources, M.M; Supervision., M.M; Funding acquisition, M.M and M.E.K; Original draft preparation M.E.K; Review and editing, M.M. All authors contributed to the article and approved the submitted version.

## Acknowledgments

The authors would like to thank Prof. Nigel Temperton for his suggestions regarding the entry model for RABV in different species and Dr. Lucy Jackson-Jones for her guidance on flow cytometry gating analysis.

## Conflict of Interest

The authors declare that the research was conducted in the absence of any commercial or financial relationships that could be construed as a potential conflict of interest.

## Notes

### Competing Interest Statement

The authors have declared no competing interest.

